# A MICROFLUIDIC DEVICE FOR LONG-TERM MAINTENANCE OF ORGANOTYPIC LIVER CULTURES

**DOI:** 10.1101/2022.06.18.496606

**Authors:** José M. de Hoyos Vega, Hye Jin Hong, Kevin Loutherback, Gulnaz Stybayeva, Alexander Revzin

## Abstract

Liver cultures may be used for modeling disease progression, testing therapies and predicting drug induced liver injury. The complexity of the liver cultures has evolved over the years from monocultures of hepatocytes to co-cultures with non-parenchymal cells and finally to precision cut liver slices. The latter culture format retains biomolecular and cellular complexity of the native liver and therefore holds considerable promise for in vitro testing. However, liver slices remain functional for ~72 h in vitro and hold only limited utility for some of the disease modeling and therapy testing applications that require longer culture times. This paper describes a microfluidic device for longer-term maintenance of functional organotypic liver cultures. Our microfluidic culture system was designed to enable direct injection of liver tissue into a culture chamber through a valve-enabled side port. Liver tissue was embedded in collagen and remained functional for up to 31 days as highlighted by continued production of albumin. These organotypic cultures also produced bile and urea and expressed several enzymes involved in metabolism of xenobiotics. In contrast, matched liver tissue embedded in collagen and cultured in the same media in a 96-well plate lost its phenotype and function on the timescale of 3 to 5 days. The microfluidic organotypic liver cultures described here represent a significant advance in liver cultivation and may be used in the future for modeling liver diseases or for individualized selection of liver-directed therapies.

## INTRODUCTION

The liver is the metabolic center of the body and is often impaired by infections or toxicants including drugs and alcohol. There is a long-standing interest in establishing in vitro liver cultures for modeling liver diseases, evaluating drug induced liver injury (DILI) and testing liver-directed therapies.^1^ Given that the majority of liver functions including energy metabolism, breakdown of xenobiotics, synthesis of serum proteins, and production of bile and urea, is carried out by hepatocytes, there has been a long-standing focus on culturing these cells in vitro.

Hepatocytes are highly differentiated epithelial cells that rapidly lose their phenotype and liver function when plated in vitro.^2–4^ Studies published over the past 30 years have systematically explored reintroduction of cellular and biomolecular elements of the liver microenvironment in vitro to enhance hepatic function. An early example of this was the development of collagen gel sandwich cultures which extended maintenance of hepatic function to 14 days and beyond^5^. Another example was the development of co-cultures, both random and micropatterned, where function of hepatocytes was rescued and maintained by the proximity of non-parenchymal cell (typically but not always fibroblasts).^6,7^ Other efforts have focused on 3D or spheroid cultures of hepatocytes alone^8,9^ and together with nonparenchymal cells^10,11^. The drive to increase complexity of hepatic cultures has culminated in precision cut liver slices which have the advantage of retaining both cellular and molecular elements of the in vivo microenvironment. However, liver slices reported to date remain functional for ~3 to 5 days and are not well suited for longer term disease modeling or drug testing experiments^12^. The objective of our paper was to establish and characterize microfluidic organotypic liver cultures that maintain hepatic function for >7 days.

Microfluidic devices are increasingly being used for cultivation of cells in general and liver cells in particular.^4^ The benefits of microfluidic devices include the ability to precisely tune chemical and mechanical stimulation experienced by the cells. In the context of liver cultures, there has been considerable interest in leveraging microfluidic devices to recapitulate liver zonation, gradients of oxygen and chemicals, that exist along the periportal to perivenous axis in the liver.^4,13,14^ Additional studies have mimicked sinusoidal architecture of the liver both in terms of geometry^15,16^ and cellular composition.^17^ There have also been some reports of organotypic liver cultures in microfluidic devices.^18–22^ These were PDMS devices that resembled a multi-well plate in that pieces of liver could be placed from the top into wells but also contained a network of microfluidic channels that delivered media and nutrients from the bottom.^23^ Despite their considerable promise and multiple advantages, hepatic function in these microfluidic multi-well plates was only assessed on the timescale of 72 hr. Therefore, to the best of our knowledge, longer-term (>7 days) function of organotypic liver cultures has not been demonstrated to date either in microfluidic or standard cultures.

Beyond the possibilities for controlling flow and establishing gradients, microfluidic devices have another interesting feature in that a relatively large number of cells may be confined to a small local volume inside a microfluidic channel. Early reports from Beebe and Levchenko labs predicted that microfluidic cell cultures operating under diffusion-dominant transport regime (minimal or no flow) should experience greater accumulation of autocrine/paracrine signals compared to standard (large volume) cultures.^24–26^ Later, presence of such autocrine signals was confirmed by other researchers who demonstrated that phenotype of stem cells cultured in a microfluidic device was affected by frequency of volume exchanges.^27^ This confirmed, albeit indirectly, importance of autocrine/paracrine signals accumulating in microfluidic devices.

Our team has also been interested in the effects of microfluidic confinement on a range of cell types.^11, 28–30^ Particularly striking to us were the observations that primary rat hepatocytes maintained epithelial phenotype and hepatic function for over three weeks as monolayer cultures in a microfluidic device.^31^ This was in stark contrast to de-differentiation and loss of hepatic function that occurs over the course of 5 to 7 days in monolayer hepatocyte cultures in standard (large volume) cultures. We determined that phenotype enhancement in microfluidic cultures was associated with the upregulated production and signaling of several hepato-inductive growth factors of which hepatocyte growth factor (HGF) was most prominently expressed. More recently, we demonstrated that similar phenotype enhancement was observed for hepatocyte spheroids cultured in microfluidic devices.^32^

In the present paper, we wanted to test a hypothesis that microfluidic confinement will result in improved phenotype and function of organotypic liver cultures. To test this hypothesis, we developed a microfluidic device (see **Figure 1**) which contained a side port for injecting a needle core of freshly harvestedliver tissue directly into the culture chamber. High levels of hepatic function persisted in this microfluidic device over the course of 30 days.

**Figure 1:**
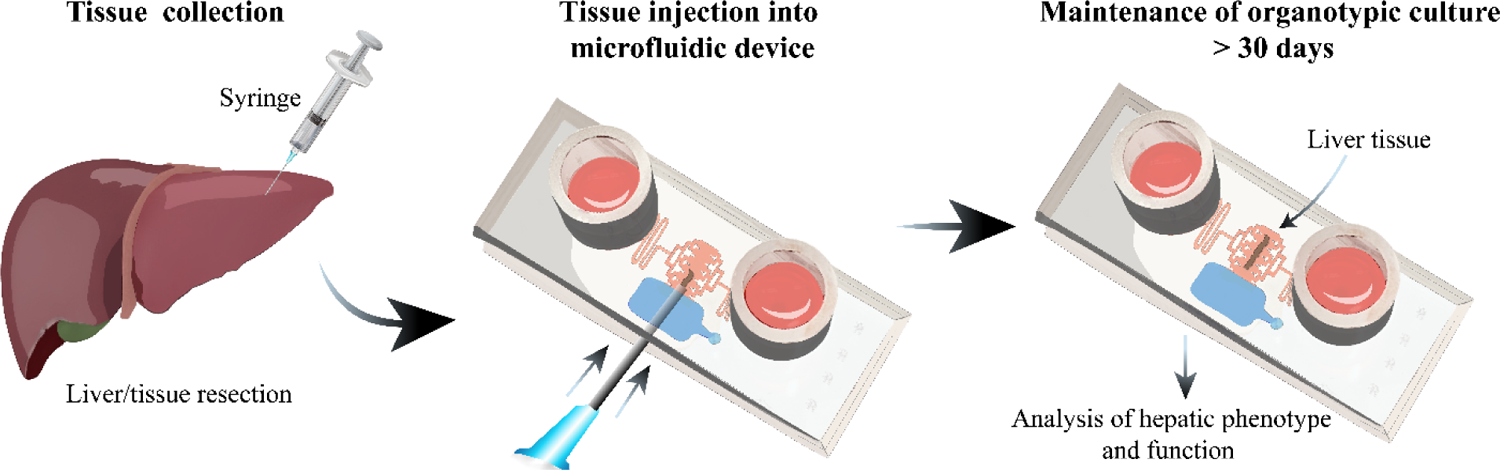
A microfluidic device for cultivation of liver tissue. Rat or human liver tissue was harvested using a 23 Ga needle and then inserted into the culture chamber of the microfluidic device. These organotypic liver cultures remained functional for up to 30 days.

## MATERIALS AND METHODS

### Reagents

The following reagents were purchased from Sigma-Aldrich: propylene glycol monomethyl ether acetate (PGMEA, 484431), chlorotrimethylsilane (386529), Pluronic F-127 (P2443), Glucagon (G2044), hydrocortisone sodium succinate (H2270), and Tween 20 (P9416). The following reagents were purchased from Thermo Fischer Scientific: Dublecco’s Modified Eagle Medium (DMEM, MT-10-013-CV), penicillin-streptomycin (15140122), Gibco Fetal Bovine Serum (FBS, 10437028), Casein (37582), Quant-iT™ PicoGreen® dsDNA Assay Kit (P7581), and Urea assay kit (DIUR-100). Epidermal Growth Factor (EGF, 3214-EG-100) and Cultrex Rat collagen I, lower viscosity (3443-100-01) were purchased from R&D Systems. HGF (SU11274) and TGFβ-1 (A83-01) inhibitors were purchased from Cayman Chemical and StemCell technologies, respectively. EZScreen TM Alanine Aminotransferase (ALT or SGPT) Activity assay kit (K941), UGT activity assay/ligand screening kit (K692), and PicoProbeTM LDH-Cytotoxicity Fluorimetric assay kit (K314) were purchased from BioVision. Albumin ELISA kit (E110-125,) and Total Bile acid assay kit (MET-5005) where purchased from Bethyl Lab and Cell Biolabs, respectively. P450-Glo CYP1A2 induction/inhibition assay system (V8771) was purchased from Promega. Sylgard 184 poly(dimethylsiloxane) (PDMS) kit (2065622) was purchased from Ellsworth and SU-8 2100 from Kayaku Advanced Materials. Insulin (Novolin N, 100 units/mL) and Ketamine (Ketalar, NDC 42023-115-10) were purchased from Novo Nordisk and Par Pharmaceutical, respectively.

### Fabrication of microfluidic devices

The valve and flow master molds were fabricated using standard photolithography techniques.^24^ Briefly, two 4-inch silicon wafers (452, University Wafer MS) were cleaned by exposure oxygen plasma (YES-G500, Yield Engineering Systems, Inc.) at 30 watts for 10 min and spin coated with SU-8 2100 to obtain a thickness of 300 μm (WS-650-23, Laurell Technologies Corporation). The wafers were prebaked at 95 °C for 1 hr as recommended by the manufacturer. A Micropattern Generator (μPG 101, Heidelberg Instruments) was used to expose the SU-8 2000 layer at a time exposure of 370 mJ/cm^2^. After a post-bake 30 step, the wafers were placed into the PGMEA developer solution for 5-10 min, followed by a hardbake step of 2 h at 135 °C. Then the wafers were treated for 2 h with chlorotrimethylsilane to minimize stiction of PDMS replicas.

A microfluidic device was fabricated using multiple layers of PDMS as described in **Figure S1**. The device was composed of four layers of PDMS that can be divided into two mirrored assemblies. The top and bottom assemblies consisted of a flow layer that contained a culture chamber, transport channels, and the injection port; while the control layer had a valve located on the injection port overlapping area. A 5:1 and 20:1 weight ratio of PDMS elastomer and curing agent were poured on the control and flow master molds, respectively, and partially cured at 80 °C for 18 min. After punching the 0.5mm in diameter inlet of the designated valve of the top assembly, the partially cured PDMS control layers were aligned onto the flow layer and baked for 2h at 80 °C. Next, 2mm diameter inlets were punched for media reservoirs and another 0.5 mm diameter hole was made for applying vacuum to a valve. Then the valve of the bottom assembly was partially punched. A strip of invisible tape approximately 2mm x 7mm was placed along the needle injection port of both top and bottom assemblies to protect the PDMS from the plasma treatment. Afterwards, the top and bottom assemblies were exposed to oxygen plasma at 30 watts for 3 min. The tape strips were then removed, the top and bottom assemblies were aligned and bonded. Two PYREX cloning cylinders (8mm (d) x 8 mm(h)) were secured at inlet and outlet dispensing 20 μL of PDMS prepolymer and curing at 80 °C for 30 minutes. Subsequently, the devices were incubated with 1% Pluronic in 1x PBS at 4°C overnight.

### Collecting, injecting, and maintaining rat liver tissue in microfluidic devices

All animal experiments were performed under the National Institutes of Health (NIH) guidelines for ethical care and use of laboratory animals with the approval of the Institutional Animal Care and Use Committee (IACUC) of the Mayo Clinic, Rochester, MN. The liver was carefully dissected from adult female Lewis rats (weighing 90–200 g, Charles River Laboratories, Boston, MA) after exsanguination with Ketamine sedation in a biosafety hood type II. The liver was then quickly immersed 3 times in cold Krebs Ringer buffer (KRB) solution in a 100mm x 25mm Petri dish to remove blood and afterwards placed in a new Petri dish containing cold KRB. A 1ml BD Luer-Lok syringe (309628, BD) with 23Ga needle was used to punch needle cores of ~ 350 μm in diameter and 3-4 mm in length.

The devices were prepared for liver tissue cultivation in the following way. First, devices were flushed with 1xPBS at 4°C for 10 min to remove Pluronic, then they were filled with hepatocyte culture media containing 10% collagen I. This media, called CH media for complete with hormones, was based on DMEM and was supplemented with 1% (v/v) penicillin-streptomycin, 10% FBS, 0.5 U/mL Insulin, 20 ng/mL EGF, 7 ng/mL glucagon and 7.5 μg/mL hydrocortisone. Devices were kept at 4°C prior to use to ensure even distribution of the collagen solution and prevent its premature gelation.

Prior to tissue culture, a device was taken out of the freezer and connected to a vacuum line inside a biosafety hood. The negative pressure from a vacuum line was used to actuate the normally closed valve and open the injection port to gain access to the culture chamber. Once the valve was actuated, the needle with tissue was placed into the injection port and transferred into the cell culture chamber by gently pressing on the syringe plunger. With liver tissue inside, vacuum was disconnected, the valve reverted to its normally closed state and the device was incubated for 2h at 37°C in CH media with 10% collagen to promote collagen gelation. Afterwards, collagen-containing media was removed and replaced by 300 μL of fresh CH media (150 μL per reservoir). The media was aspirated from the reservoirs and exchanged every 24 hr.

In parallel with microfluidic organotypic cultures, we also created conventional (control) cultures collected from the same livers using the same syringes and needles. These needle cores were plated into wells of a 96-well plate (25-109, Genesee Scientific) pretreated with 1% Pluronic in 1x PBS buffer as described above. First, 50 μl of CH media with 10% collagen were dispensed in the well and incubated at 4°C for 10-15 min. Then the plate was placed in the biosafety hood and liver cores were dispensed into each well. The well plate was incubated for 2h at 37°C to allow for collagen gel formation. Afterwards, 250 μl of fresh CH media without collagen was added into the wells. Media in these well-based cultures was changed every 24h for the duration of the experiment.

### Obtaining and culturing human liver tissue

Normal adjacent and diseased human resection samples were obtained from adult patients undergoing surgical resection at Mayo Clinic Rochester using IRB protocol #17-010608. Per IRB protocol, these were pieces of liver < 1 cm^2^ in area. Resected tissue was collected into 50 mL conical tubes filled with University of Wisconsin (UW) cold storage solution for transportation and storage prior to punching of needle cores. The samples in UW were placed in a 6mm Petri dish and needle cores were collected and transferred into microfluidic devices and wells of a 96-well plate as described above for rodent tissue. CH media with 10% collagen I solution was used for embedding liver cores in collagen gel. Subsequently, liver tissue was cultured in CH media without collagen for the duration of a multiday experiment.

### Analysis of hepatic function

Assays of albumin, LDH, ALT, total bile acids, urea, CYP1A2, and UGT were carried out in accordance with manufacturer’s instructions. For CYP1A2 induction and UGT activity, samples were treated with 100μM omeprazole and 100μM Bilirubin for 24 hr, respectively, prior to tissue collection. Optical density measurements were made using UV/vis spectrophotometer (Synergy H1, BioTek).

For cell number quantification, liver needle cores were collected and placed individually in 1.5 mL Eppendorf–s tubes. Tissue was lysed using 20mM Tris-HCl, 2mM EDTA and 1% Tween 20 at pH 6.9 and vortexed for 10 min. Picogreen assay kit was used per manufacturer’s instructions. Optical density measurements were made using UV/vis spectrophotometer.

### Characterization of hepatic gene expression by RT-PCR

Tissue samples at day 0 and 7 were collected from devices and dissociated in the lysis buffer (Qiagen) by agitating with an electronic pestle for 1 min. Then lysed samples were processed with total RNA isolation kit (Roche) according to the manufacturer’s instructions. The concentration of the isolated RNA was quantified using NanoDrop One Spectrophotometer (Thermo Fisher Scientific), then complementary DNA was synthesized using a commercial kit (Roche) following manufacturer’s instructions. Gene expression level was quantified using SYBR Green (Roche) with QuantStudioTM 5 System (Thermo Fisher Scientific). All target primer sequences were purchased from IDT (See supplementary table 1). Real-time PCR was performed with 40 cycles according to the following amplification protocol: denaturation at 95□ for 5 s, annealing at 55□ for 15 s, and extension at 69□ for 20 s. The final gene expression was normalized to glyceraldehyde 3-phosphate dehydrogenase (GAPDH) through ΔΔCT method.

### Immunofluorescence staining and imaging of liver tissue

Liver cores were fixed by injecting 300 μL of 4% PFA into the device and incubating for 24 h at 4 °C. Then, a tissue was carefully removed from the device and placed into solution of 1x PBS + 30% sucrose for 1hr. Subsequently, a tissue was embedded in optimal cutting temperature (OCT) compound (Tissue-Tek), frozen and sliced using a cryostat (Leica Biosystems, CM1950, USA) to the thickness of 12 μm per slice. Tissue sections were washed with PBS, permeabilized with 0.05% Tween 20 for 30 min and blocked with casein blocking solution for 10 min. Slices were incubated overnight at 4°C with the following primary antibodies: sheep anti-rat albumin (1:100; Bethyl lab Inc.) mouse anti-multidrug resistance associated protein (MRP)-2 (1:100; Novus). Afterwards, the slices were washed with PBS and incubated for 1 h with following secondary antibodies: Alexa-488 donkey anti-sheep IgG (1:1000), Alexa-647 donkey anti-rabbit IgG, (all secondary antibodies were diluted at 1:1000). The sections were mounted with Vectashield containing 4,6-diamidino-2-phenylindole (DAPI) for cell nuclei staining. Micrographs were taken using an inverted fluorescence microscope (IX-83, Olympus) using 10x and 20x long distance objectives. Tissue size was measured using Olympus imaging software LCmicro 2.2.

### COMSOL modeling of HGF accumulation in a microfluidic device

Numerical simulations were performed with COMSOL (Burlington, MA) to model accumulation of HGF from a needle core biopsy (R=175 μm, L = 3000 μm) with ~4000 secreting cells is accumulated in the microfluidic device and a well of a 96-well plate. Assumptions made for modeling included: a constant secretion rate of 0.4 ng·h^−1^·cell^−1^, diffusion coefficient of 8.5×10^−11^ m·s^−1^ and no flow^28^. HGF concentrations in the device and microwell were generated every 10 min for 24 h.

### Statistical analysis

Statistical significance was determined using t-test analysis with p< 0.05. Typically, three or more biological replicates (n>3) were analyzed and averaged for each experimental group. GraphPad Prism (ver. 7; GraphPad Software) was used to statistically analyze the data.

## RESULTS AND DISCUSSION

### Device design and fabrication

We were motivated by several considerations in designing a microfluidic device for organotypic liver cultures. First, we wanted to test a hypothesis that placing intact liver tissue into confined volume of a microfluidic device will result in hepatic phenotype and function that would extend beyond the current state of the art of 48 to 72 hr. Second, we wanted to address the need for a culture platform that will work with minimal amounts of tissue (e.g., biopsies, small resections, fine needle aspirates) where precision slicing is not practicable. To achieve this, we wanted to design a system that may be interfaced with a syringe for transferring small amount of tissue from a needle into a microfluidic culture chamber.

The device developed by us is shown in **Figure 2A**. It is composed of PDMS and contains a flow layer sandwiched between two control layers. The flow layer contains a culture chamber connected to media reservoirs (cloning cylinders) via branching transport channels. The branching channels were implemented to ensure uniform delivery of nutrients along the long axis of the culture chamber (5 mm, see **Figure 2**). The culture chamber was designed to receive cores generated from a 23 Ga needle with typical dimensions of 2-4 mm in length and 300-400 μm in diameter. The key feature of this device was an injection port controlled by a normally closed valve. This valve was opened by applying negative pressure and 23 Ga needle was manually inserted directly into the cell culture chamber (see **Figure 2B** for a sequence of steps involved in device operation). The injection port and the culture chamber were made sufficiently tall (600 μm) to accommodate a syringe needle. Importantly, after the needle was retracted and negative pressure turned off, the valve reverted to its normally closed state, the injection port was sealed, and the device could be handled with ease in a manner similar to a Petri dish. No leakage of the media in the region of the injection port was observed over the timescale of an experiment (up to 31 days). **Video 1** highlights ease of device operation and placement of tissue into the culture chamber. **Video 2** demonstrates that the culture chamber may be accessed repeatedly with a syringe to deposit another tissue or to retrieve the original tissue.

**Figure 2:**
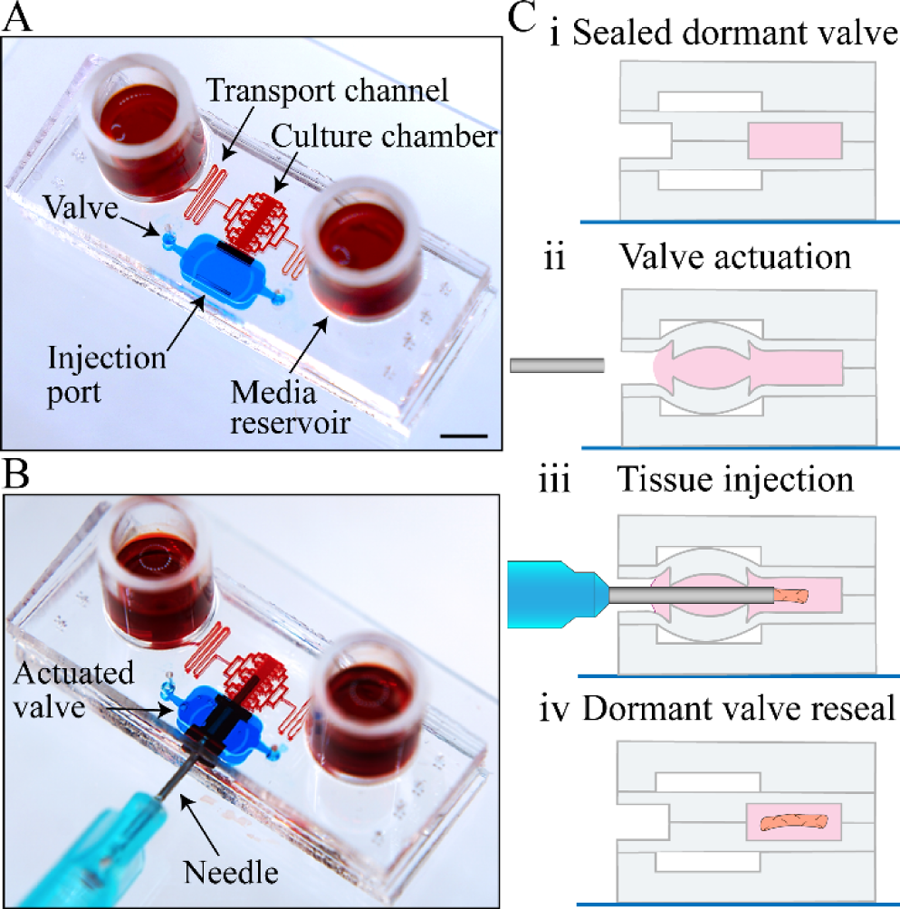
Introduction of liver needle cores into a microfluidic device. (**A**) Elements of a microfluidic device highlighted using dyes of different colors. Red dye denotes the flow layer containing culture chamber, media reservoirs and transport channels. Blue dye shows location of the injection port and the valve. Note that the valve is normally closed and that red and blue dye in the culture chamber and injection port respectively do not mix. Scale bar: 5mm (**B**) Actuation of the valve opens access to the culture chamber and allows introduction of the needle and transfer of tissue. (**C**) Cross-sectional view of the device describing steps in device operation: (i and ii) actuating the valve, (iii) needle insertion and transfer of tissue, (iv) retraction of the needle followed by return of the valve into normally closed state.

### Assessing viability and function of organotypic rat liver cultures in microfluidic devices

Given the scarcity and variable quality of human liver tissue, we relied on rat liver tissue for device validation and characterization. In order to mimic a clinical scenario, we excised rat liver and placed it in cold KRB solution. Cores were then collected by inserting a syringe with 23 Ga needle into the liver tissue. At the early stages of this study, we determined that presence of collagen gel improved functionality of liver tissue both in microfluidic devices and multi-well (data not shown). This result was unsurprising given a number of reports describing that presence of collagen or other ECM gels enhanced function of primary hepatocytes.^5,33,34^ Therefore, all of the subsequent experiments were carried out with liver tissue embedded in collagen type I gel. Another experimental consideration was the size of the liver tissue to be cultured. Given the evidence for nutrient and oxygen diffusion being limited to ~400 μm^35^, we wanted minimal dimension of the liver tissue to be <400 μm. This design parameter was satisfied by using a 23 Ga needle that produced roughly cylindrical cores with an average diameter of ~300 μm. Liver cores were placed into two types of cultures systems: 1) microfluidic devices described in **Fig. 2A** and 2) wells of a 96 well plate. The cores cultured in these two systems came from the same liver sample, were imbedded in collagen gel, and exposed to the same media (CH media, please see Methods for composition). The same volume (~300 μL) was used for both well-based and microfluidic liver cultures.

**Figure 3A-B** shows examples of liver tissue placed into microfluidic devices and microwells, respectively. Presence of collagen gel in the cultures is highlighted by Picrosirius red staining (See **Figure 2C**). Cytotoxicity in liver cultures was assessed using ALT assay, a clinical indicator of liver injury. As seen from **Figure 3D**, microfluidic liver cultures were associated with significantly lower levels of ALT in the culture media compared to well-based cultures.

**Figure 3:**
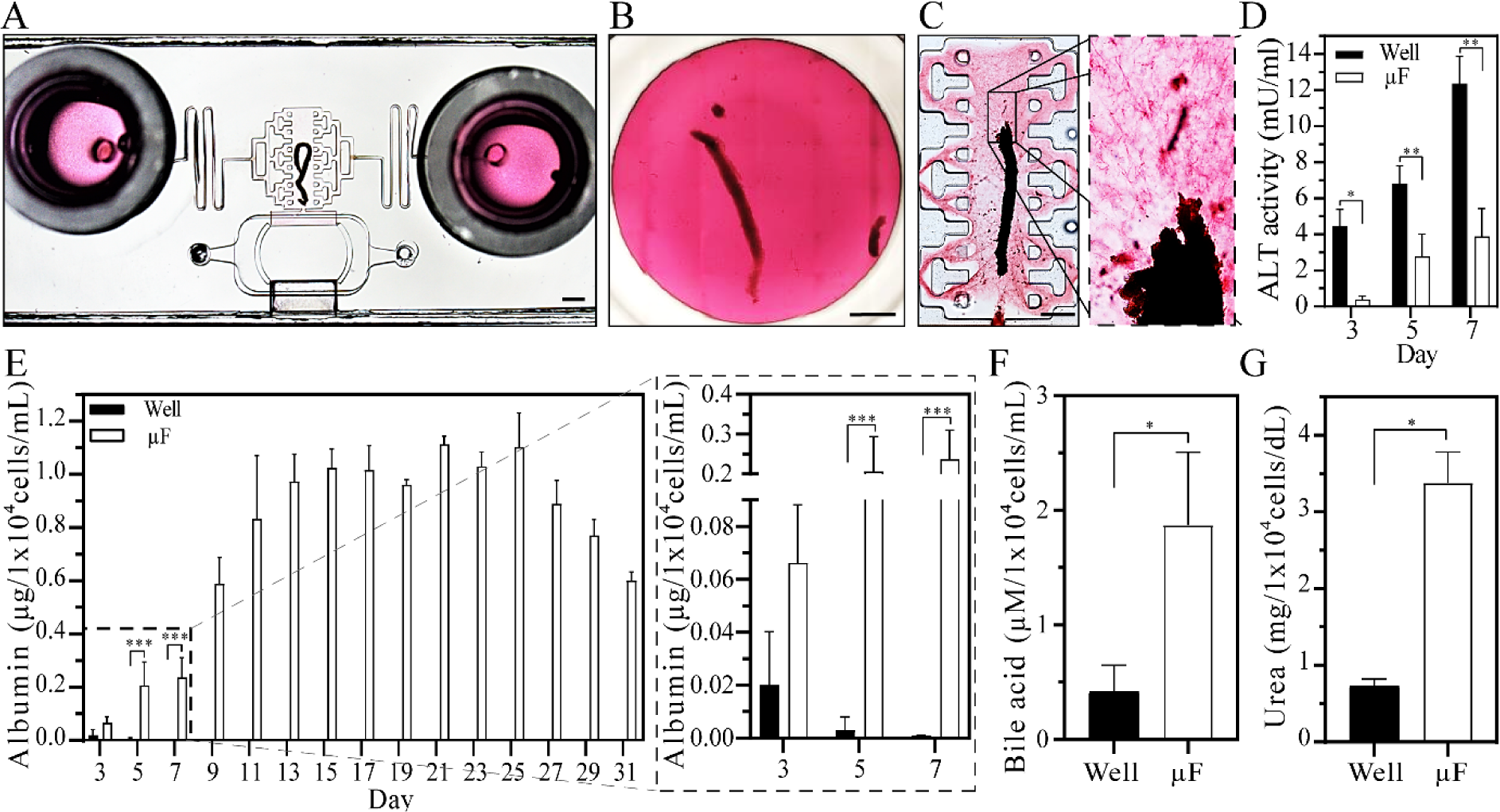
Assessing cytotoxicity and functionality of organotypic microfluidic liver cultures. (**A**) A microfluidic device with a liver core. Scale bar: 1mm. (**B**) A well of a 96 well-plate with a liver core. Scale bar 1 mm. (**C**) Picrosirius red staining highlighting presence of collagen in a culture chamber of a microfluidic device. Scale bar 1mm. (**D**) ALT activity assay to measure hepatotoxicity in standard and microfluidic cultures. Note that higher levels of ALT indicate more injury and hepatocyte damage. (**E**) Albumin synthesis of organotypic live cultures comparing well and microfluidic (μF) culture conditions. (**F**) Bile acid production for well and microfluidic liver cultures. (**G**) Urea analysis for well and microfluidic liver cultures. Data are represented as means ±SD of 3 tissue samples cultured in well and microfluidic platform. **Statistical significance** determined by two-tailed unpaired t-test, * p-value <0.05, ** p-value <0.01, and *** p-value < 0.001.

Importantly, liver tissue was not only more viable but functional in microfluidic devices. This was confirmed by albumin, urea and bile acid assays shown in **Figure 3(E-G)**. Serum albumin is produced by the liver and its synthesis in vitro is an important indicator of hepatic function. Urea is a byproduct of protein synthesis and is another frequently used indicator of hepatic function in vitro. Finally, bile acid synthesis and secretion is another important liver function involved with absorption and metabolism of nutrients in the gut.^36^ As may be appreciated from **Figure 3E**, albumin was robustly synthesized in microfluidic liver cultures over the course of 31 days. For comparison, albumin production of liver tissue in standard cultures declined precipitously in a matter of days and was undetectable beyond 7 days. At day 7, microfluidic organotypic cultures produced ~250 times more albumin compared to well-based liver cultures. In addition, bile acid was ~4 fold higher while urea was ~5 fold higher in microfluidic cultures compared to standard well-based cultures. Urea production in microfluidic cultures was sustained for 31 days with the highest point at day 21 (See **Fig. S2**). PicoGreen assay was used to determine the number of cells per liver core (~3 × 10^4^) and then convert hepatic function to per cell basis.

### Evaluating hepatic phenotype in microfluidic liver cultures

As noted earlier in this paper, liver parenchyma is comprised of hepatocytes – highly differentiated epithelial cells that perform multiple functions including synthesis of albumin and production of bile. Hepatocytes possess complex polarity. Their basolateral domains contain machinery for secreting proteins into blood vessels (sinusoids) while apical domains form channels (canals) for bile transport.^3,37^ We used immunofluorescence staining at day 7 to assess hepatic phenotype and polarization in microfluidic and well-based organotypic liver cultures. Freshly isolated rat liver tissue was also immunostained and used as the gold standard for benchmarking the two culture systems. **Figure 5A** highlights the results of immunofluorescence staining for albumin and multi-drug resistant protein (MRP-2) for three liver tissue types mentioned above. While albumin is used to assess protein synthesis capacity of hepatocytes, MRP-2 is a bile acid transporter that localizes to the apical domains and is associated with polarized and well-differentiated hepatocytes. **Figure 5B** highlights the fact that after 7 days of culture in a microfluidic device, rat liver tissue stained strongly for albumin and possessed extensive network of bile canals as indicated by MRP-2 staining. Cytoarchitecture of microfluidic liver cultures resembles that of a freshly isolated liver tissue (See **Figure 5A**). Conversely, standard (well-based) organotypic cultures had minimal expression of albumin or MRP-2 at day 7(See **Figure 5C**). The immunofluorescence results are consistent with functional analysis data presented in **Figure 3** that demonstrated several fold higher levels for albumin, bile acid and urea production in microfluidic cultures.

**Figure 5:**
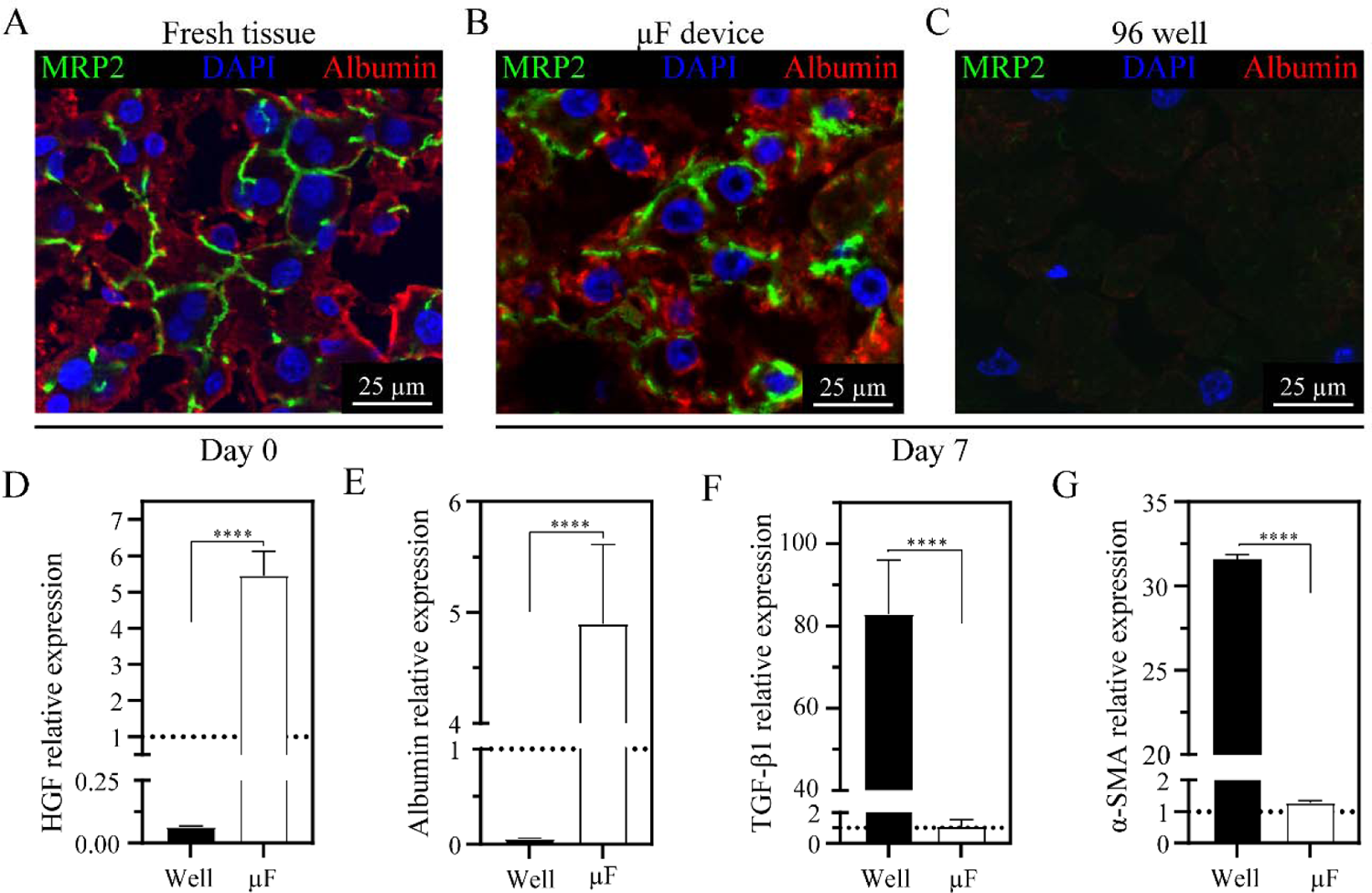
Evaluating hepatic phenotype in microfluidic organotypic cultures. Immunofluorescence staining for albumin (green) and MRP-2 (red). Comparing cytoarchitecture of freshly isolated rat liver tissue (**A**) to microfluidic (**B**) and standard cultures (**C**) at day 7. RT-PCR analysis of key genes associated with maintenance or loss of hepatic phenotype. High levels of HGF (**D**) and albumin (**E**) expression, along with low levels of TGF-β1 (**F**) and α-SMA (**G**) are associated with maintenance of hepatic phenotype. **Statistical methods:** Data are presented as means ±SD of 3 tissue samples cultured in wells and microfluidic devices. Statistical significance determined by two-tailed unpaired t-test, ****p-value < 0.0001.

Beyond immunofluorescence, we wanted to assess key genes associated with liver phenotype and function. Our prior studies with hepatocytes revealed that, when compared to standard (large volume) cultures, microfluidic cultures upregulated expression of HGF and downregulated expression of TGF-β1.^11,32^ RT-PCR analysis confirmed that similar observation held true for organotypic liver cultures. Microfluidic liver cultures had ~83-fold higher HGF expression compared to well-based cultures (see **Figure 5D**). The hepato-inductive effects of HGF are reflected in albumin gene expression which is 82-fold higher in the device compared to well-based culture format at day 7 (See **Figure 5E**). Conversely, TGF-β1 expression was 75.5-fold lower in microfluidic cultures. Interestingly, the level of TGF-β1 expression in microfluidic cultures at day 7 was similar to that of freshly isolated liver tissue (dashed line in a **Figure 5F**). This observation was highly encouraging given the negative impact of TGF-β1 signaling on hepatic function via liver fibrogenesis, activation of stellate cells and de-differentiation of hepatocytes.^38,39^ Alpha smooth muscle actin (α-SMA) is another marker of stellate cell activation and fibrogenesis in the liver. Expression of this marker in microfluidic liver cultures at day 7 was comparable to α-SMA levels in freshly isolated liver tissue (See **Figure 5G**). Conversely, well-based liver tissue expressed α-SMA at levels 24-fold higher than microfluidic cultures at day 7. This gene expression analysis was consistent with microscopy-based observations of proliferating stromal cells in standard (well-based) cultures (data not shown).

### Metabolism of xenobiotics in microfluidic liver cultures

Liver functions as the metabolic center of the body where xenobiotics are modified by hepatic enzymes. Such enzymes are expressed by the hepatocytes and are categorized as phase I and phase II.^40^ Cytochrome p450 enzymes (CYPs) are largely responsible for phase I metabolism of the liver.^41–43^ The expression and activity of these enzymes decrease precipitously as hepatocyte de-differentiate in vitro. We carried out RT-PCR analysis for three of the more common CYPs: CYP1A1, CYP1A2, and CYP2E1. These are phase 1 enzymes responsible for metabolizing a wide range of compounds including melatonin, caffeine, and ethanol.^41–43^ As seen from **Figure 6**, CYP expression in microfluidic liver cultures was manyfold higher (37 to 260 depending on the enzyme) than in well-based cultures at day 7. It was encouraging to observe that levels of CYP1A2 expression in microfluidic cultures at day 7 were comparable to those of freshly isolated rat liver tissue (**Figure 6A**). Interestingly, CYP1A1 expression was higher in microfluidic cultures compared to freshly isolated tissue (**Figure 6B**). This may be attributed to supplementation of media with hydrocortisone which is metabolized by CYP1A1 and likely induces expression of this enzyme.^44,45^ While ethanol metabolizing enzyme CYP2E1 was expressed at a lower level in a microfluidic liver culture compared to fresh liver tissue, its expression was still 260-fold higher than that of well-based liver tissue (See **Figure 6C**). In addition to characterizing levels of CYP gene expression, we assessed activity of CYP1A2 by challenging organotypic liver tissue with 100 mM omeprazole, a compound metabolized by this enzyme, at different times during culture. **Figure 6D** highlights 2-fold induction in CYP1A2 levels compared to control microfluidic cultures that did not receive omeprazole. It is also worth pointing out the similarity in induction of CYP1A2 at days 3 and 7 of culture. To the best of our knowledge maintenance of CYP1A2 expression in organotypic liver cultures over 7 days at levels similar to freshly isolated liver tissue has not been reported in the literature. This highlights, once again, the maintenance of hepatic function enabled by the microfluidic organotypic cultures.

**Figure 6:**
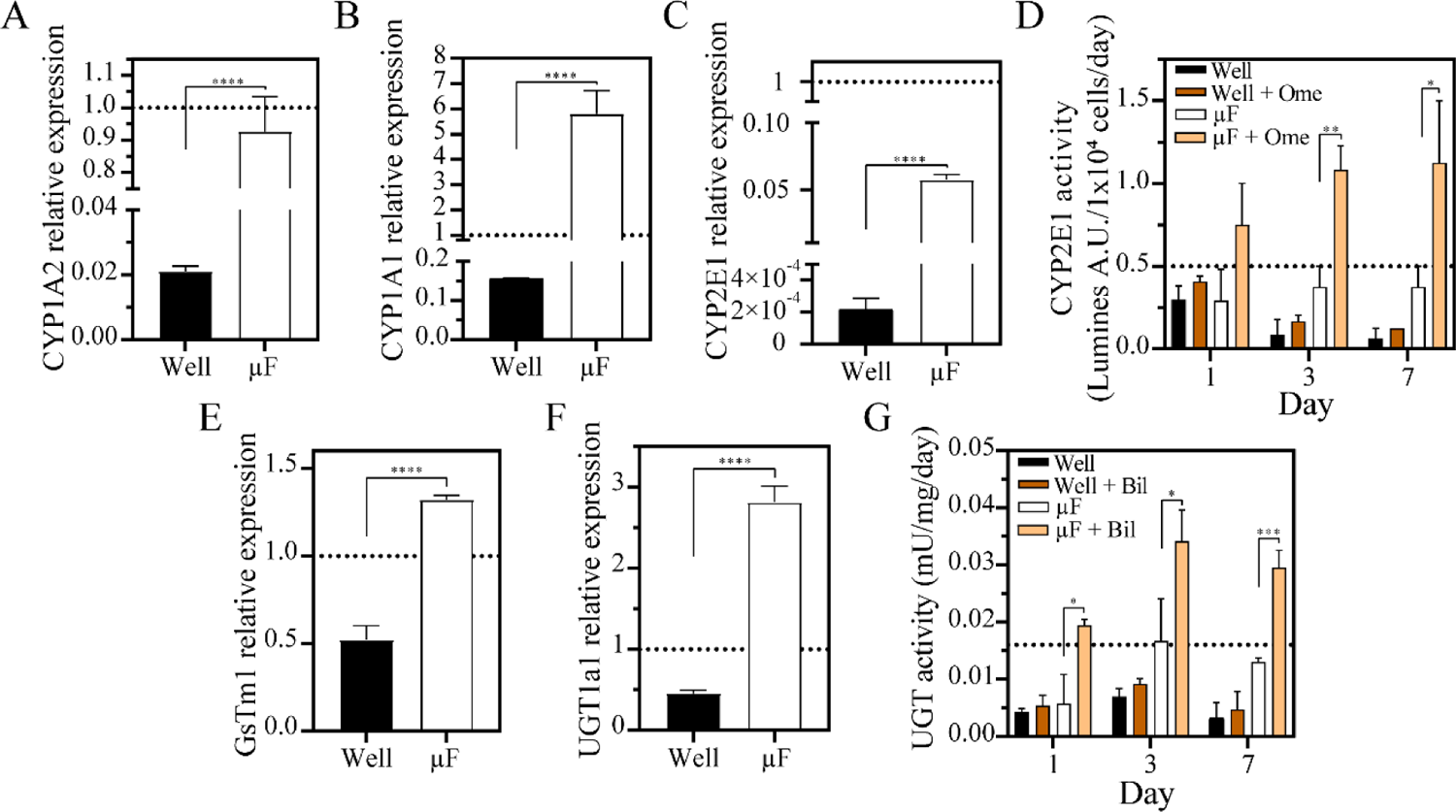
Assessing Phase I and Phase II metabolism in microfluidic liver cultures. (**A-C**) RT-PCR analysis of phase I enzymes at day 7 of culture, CYP1A2 (A), CYP1A1 (B) and CYP2E1 (C). Dashed line represents gene expression of freshly isolated liver tissue (day 0). (**D**) CYP1A2 activity was assessed by adding omeprazole (Ome) into culture media. Dashed line refers to CYP1A2 activity of freshly isolated rat liver tissue. (**E,F**) RT-PCR analysis of phase II enzymes, GST (E) and UGT (F). Dashed lines correspond to freshly isolated tissue. (**G**) UGT activity assessed using bilirubin (Bil) as a substrate. Dashed line corresponds to freshly isolated tissue. **Statistical methods:** Data are represented as means ±SD of 3 tissue samples cultured in well and microfluidic platform. Statistical significance determined by twotailed unpaired t-test, * p-value <0.05, ** p-value <0.01, ***p-value <0.001 and **** p-value < 0.0001.

The liver plays an important role in improving solubility of xenobiotics to enhance their excretion via bile or urine. This function is performed by phase II enzymes.^46^ Glutathione S-transferase (GST) and UDP-glucuronosyltransferase (UGT) are some of the more common phase II enzymes responsible for conjugating glutathione and glucuronic acid respectively onto xenobiotic compounds. RT-PCR analysis revealed that GST and UGT gene expression at day 7 of microfluidic cultures was 0.6- and 3-fold higher respectively compared to freshly isolated tissue (dashed line) (**see Figure 6E and F**). It is likely that the overexpression of these enzymes is caused by the supraphysiological levels of insulin and hydrocortisone in the media.^47–50^ Phase II enzymes in a standard culture format exhibited 6.29 fold lower expression compared to microfluidic cultures (See **Figure 6G**).

### What is the reason for improved function of microfluidic organotypic liver cultures?

As noted earlier in this paper, microfluidic confinement was previously reported by us and others to affect phenotype of multiple cell types, including hepatocytes.^26,31,32,51^ To check whether this observation held true for liver tissue, we designed an experiment so as to minimize differences between standard (well-based) and microfluidic culture formats. Liver cores were collected in identical manner, collagen gel embedding, media type and the volume (300 μL) were the same for both culture formats. The key difference between the culture systems was the distribution of media around the liver tissue. In a microfluidic device, liver tissue was placed into a small local volume of the culture chamber (2.3 μL) connected to the transport channels (2.7 μL). The cylindrical reservoirs placed at the inlet and outlet of the device contained 295 μL (**see Figure 2**). COMSOL was used to set up a model that reflected geometry of the microfluidic device and took into account secretion and diffusion of a putative signaling molecule (70 KDa, similar to HGF, see Table S1 for model parameters). As may be appreciated from **Figure 7A**, 98% of secreted signal is expected to be retained in the culture chamber, in the immediate vicinity of liver tissue. Concentration of this putative hepato-inductive signal at the surface of liver tissue is expected to reach ~37 nM in the microfluidic cell culture chamber compared to 1.48 nM in a well of a 96-well plate (see **Figure 7B**). What endogenous/autocrine signals are responsible for improved liver function in microfluidic devices? We previously carried out sequencing and identified several hepato-inductive signals to be upregulated in microfluidic cultures of hepatocytes.^31^ While these signals also included EGF, IGF and KGF, expression of HGF was most prominently upregulated by microfluidic confinement. In the present study, we also observed ~83-fold higher HGF expression in microfluidic vs. standard organotypic liver cultures (see **Figure 5A**). To further explore the importance of HGF signaling, we supplemented culture media with 5 μM SU11274 (inhibitor of HGF receptor c-met). As shown in **Figure 7C**, treatment with c-met inhibitor resulted in dramatic, 5-fold, decrease in production of albumin which confirmed important role of endogenous HGF signaling in microfluidic liver cultures. Interestingly, albumin production persisted beyond 7 days in the presence of inhibitor. This may suggest that concentration of the inhibitor was insufficient to degrade hepatic phenotype or more likely that HGF acts in concert with other secreted signals to shape hepatic phenotype in microfluidic confinement.

**Figure 7.**
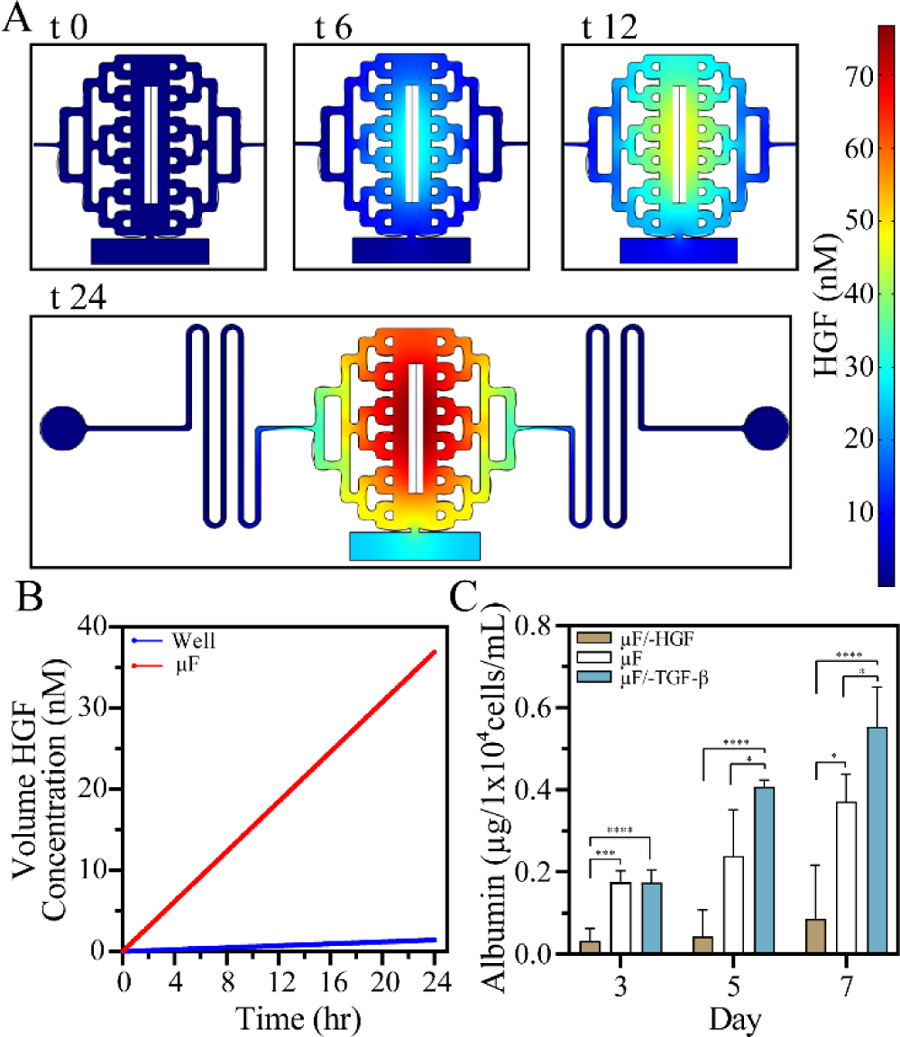
Exploring the role of endogenous signals in microfluidic liver cultures. (**A**) Modeling of HGF distribution in a microfluidic device over the course of 24 h under static conditions. (**B**) Changes in HGF concentration next to the liver tissue in a microfluidic device and a well. (**C**) Modulation of hepatic function in the microfluidic liver cultures with HGF and TGF-β1 inhibitors. **Statistical methods**: Data are presented as mean ±SD of 3 microfluidic devices. Statistical significance determined by two-tailed unpaired t-test, * p-value <0.05, *** p-value < 0.001, and **** p-value < 0.0001.

In addition to degrading hepatic function by interfering with HGF signaling, we asked whether hepatic function may be further improved by inhibiting TGF-β1 – a signal that promotes liver fibrosis and contributes to loss of hepatic phenotype. As shown in **Figure 7C**, supplementing media with TGF-β inhibitor (5 μM A83-01) resulted in modest but statistically significant enhancement in albumin production. The modeling and experimental results presented in **Figure 7** highlight that geometry of the culture system may be designed to enhance accumulation of endogenous signals and that such signals exert a powerful effect on phenotype and function of cultured liver tissue.

### Assessing the presence of nonparenchymal cells in microfluidic liver cultures

In addition to hepatocytes, parenchymal cells of the liver that comprise 80% of cell mass, the liver also contains nonparenchymal cells including stellate cells, sinusoidal endothelial cells and leukocytes. We used immunofluorescence staining and RT-PCR analysis to evaluate cellular heterogeneity of the organotypic liver cultures. Hepatic stellates cells are normally quiescent in healthy liver and acquire a fibrotic, activated phenotype during injury.^52^ Immunofluorescence staining for desmin confirmed the presence of quiescent-like stellates cells in microfluidic cultures that resemble those found in fresh tissue (see **Figure 8A,B**). In contrast, well-based liver cultures contained striated activated stellate cells at day 7 (see **Figure 8C**). Microscopy observations were supported by RT-PCR analysis of α-SMA – a marker of stellate cell activation. As may be appreciated from **Figure 5G**, α-SMA expression was significantly higher in well-based liver cultures compared to microfluidic cultures. It is also worth noting that α-SMA expression was comparable in freshly isolated tissue and in microfluidic devices after 7 days of culture. Glial fibrillary acidic protein (GFAP) is a hepatic stellate cell-specific marker. While typically associated with stellate cell quiescence, GFAP was recently shown to become upregulated at early stages of stellate cell activation.^53^ Our results indicate that GFAP expression is retained in microfluidic cultures, albeit at a significantly lower level compared to freshly isolated tissue. Taken together, microscopy observations and RT-PCR analysis of GFAP, α-SMA and TGF-β1 support the notion that hepatic stellate cells are retained in microfluidic liver cultures and that these cells maintain a more quiescent phenotype compared to well-based cultures.

**Figure 8:**
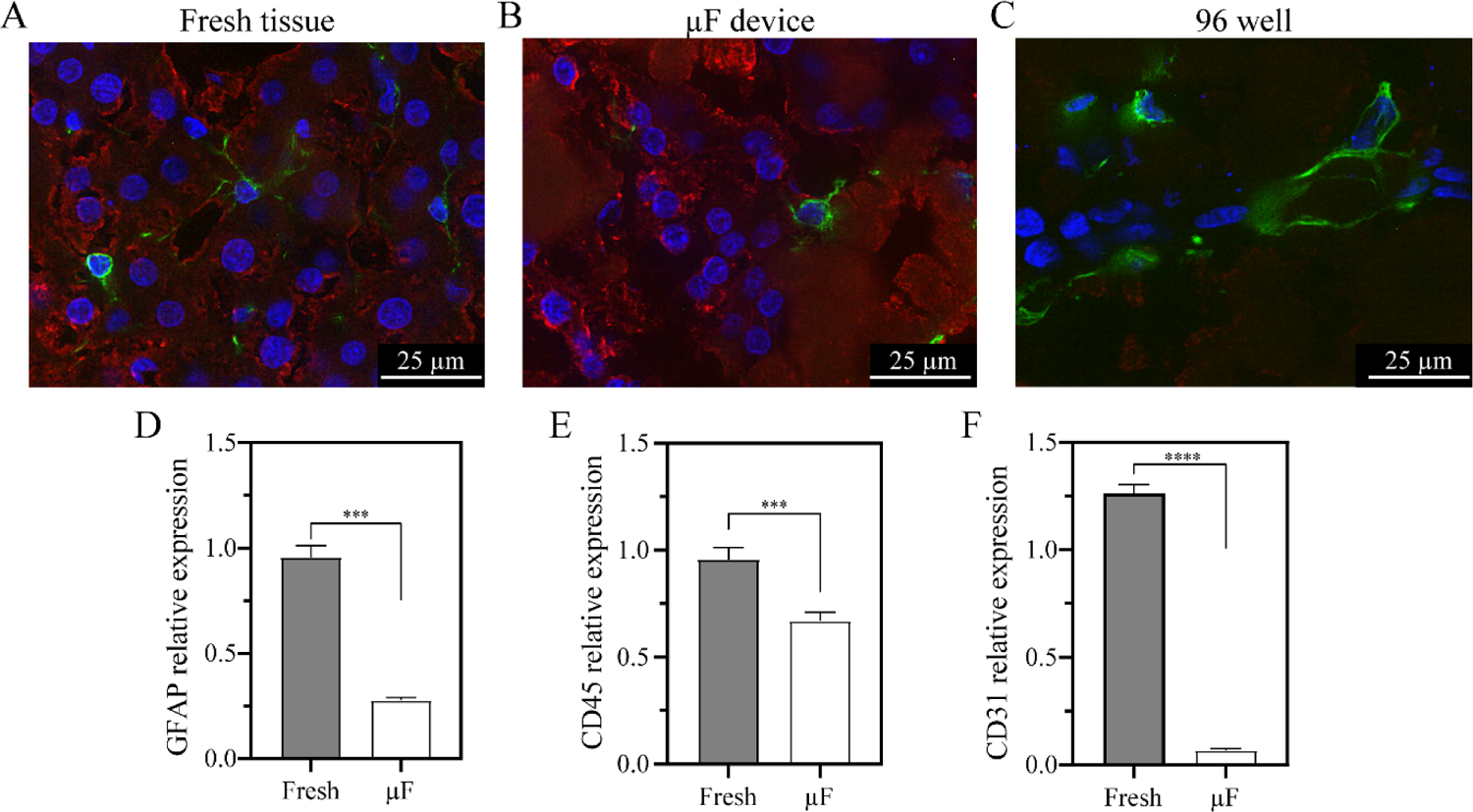
Characterization of nonparenchymal liver cells. (**A-C**) Immunofluorescence staining for desmin - a marker of hepatic stellate cells. Freshly isolated liver tissue (A) was compared to microfluidic liver tissue (μF) (B) and well-based culture (C). Note that rounded desmin-expressing cells are associated with quiescence while elongated desmin-expressing stellate cells are activated. Also note that albumin expression is not observed in well-based cultures. (**D**) RT-PCR analysis GFAP expression in fresh tissue and microfluidic cultures at day 7. (**F**) RT-PCR analysis of endothelial marker PECAM-1 (CD31) in fresh tissue and microfluidic cultures at day 7. (**E**) RT-PCR analysis of pan-leukocyte marker CD45 in freshly isolated liver tissue and microfluidic cultures at day 7. **Statistical methods:** Data are represented as means ±SD of 3 microfluidic devices. Statistical significance determined by two-tailed unpaired t-test, ** p-value <0.01, *** p-value < 0.001, and **** p-value < 0.0001.

In addition to stellate cells, we also characterized presence of leukocytes. RT-PCR analysis for pan-leukocyte marker, CD45, and immunofluorescence for CD45 and CD163 were used to assess presence of these cells. As seen from **Figure 8E**, levels of CD45 expression in microfluidic cultures were only marginally lower than in freshly isolated liver tissue. Immune cells were also observed in the microfluidic tissue via microscopy (data not shown). We should note however that both RT-PCR analysis (**Figure 8F**) and immunofluorescence staining (not shown) indicated rapid loss of liver sinusoid endothelial cells in the microfluidic (and well-based) cultures. There are several reports describing endothelial cell/hepatocyte co-cultures, which means retention of endothelial cells may be achieved in the future by further refining media composition. At the present time, our microfluidic culture system rescues and maintains liver parenchymal cells and supports some but not all non-parenchymal cell types.

### Establishing microfluidic cultures of human liver tissue

The ability to culture human liver cells opens exciting opportunities for disease modeling or therapy testing; however, liver tissue samples are typically collected as small wedges from liver resections or as needle core biopsies and are not amenable to isolating functional hepatocytes. A microfluidic device described in this paper obviated the need for tissue digestion and isolating human hepatocytes and was ideally suited culturing human liver tissue. However, before proceeding to human liver cultures, we wanted to assess how to best store tissue prior to transfer into a microfluidic device. Unlike experiments with rat livers described above which were carried out in a research lab setting where tissue was cultured immediately upon liver excision, human liver resections or biopsies are collected in a clinical setting and become available for research use after delay of tens of minutes to hours. Given that liver tissue rapidly becomes ischemic after excision,^54^ we wanted to assess what preservation solution was best suited for storage tissue post procurement and how time of storage prior to cultivation affected hepatic function in microfluidic devices. Rat liver tissue was used to explore these effects. Right and left liver lobes were excised and placed into cold preservation solutions; KRB (left lobe) and UW (right lobe). Cores were collected every 1h over the course of 3h and were then placed into microfluidic devices. As seen from **Figure S3A**, liver pieces in KRB solution became whitish and necrotic at the puncture sites after 1h, with necrotic regions spreading by 3h timepoint. Conversely, liver piece is UW solution remained pale reddish without signs of necrosis (**See Figure S3B**). Albumin production was assessed for tissue stored for different durations of time in either KRB or UW solutions. As seen from **Figure S3(C,D)**, liver tissue collected after 1h storage in UW solution was most functional after 7 days of culture. Unfortunately, it appears that the advantage of UW solution was lost beyond 1h of incubation as albumin production was similar for all other conditions (UW and KRB). These results underscore the need for timely (within 1h) transfer of human liver tissue into microfluidic cultures.

Having established the benefits of using UW solution for storage of liver tissue and the need to minimize storage time, we proceeded to culture adjacent normal liver tissue from 3 patients with liver disease. The liver tissue was injected into microfluidic devices and maintained over the course of 7 days (**Figure 9A**). As demonstrated by **Figure 9 (B,C)**, human liver cultures had a constant ALT levels and maintained albumin production during this timeframe albeit at much lower levels than those described for rat liver tissue in **Figure 3D**. These differences may be explained in part by interspecies differences in albumin production and by the fact that while human liver tissue was obtained from adjacent normal regions, it did come from patients with liver disease and lower hepatic function. The results presented in **Figure 9C** are highly encouraging as they demonstrate the ability to maintain function of patient-specific liver tissue for 7 days.

**Figure 9:**
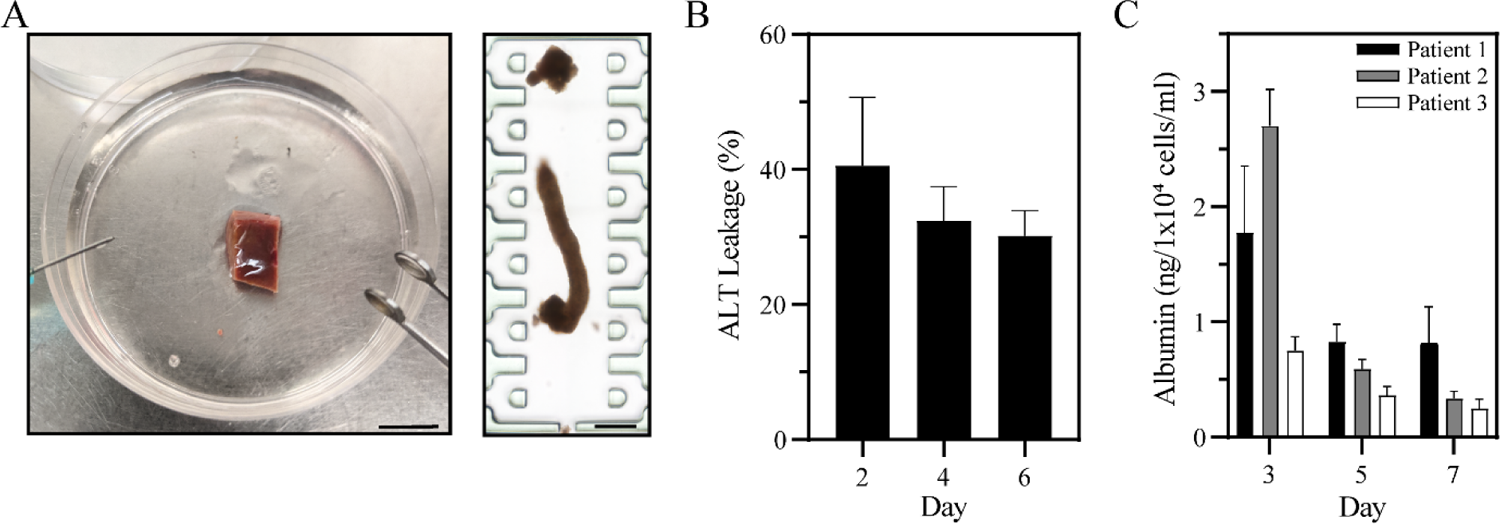
Culturing human liver tissue in microfluidic devices. (**A**) Photographs of resected liver sample in a P100 dish and needle core in a microfluidic device. Scale bar: 1 cm and 500 μm. (**B**) ALT is an enzyme released into media by dying hepatocytes. ALT assay may be used to specifically assess hepatotoxicity in organotypic cultures comprised of multiple cell types. (**C**) Production of albumin evaluated for 3 patient liver samples over the course of 7 days

### How do our microfluidic cultures compare to existing organotypic liver cultures?

Precision cut liver slices (PCLS) were first described by Otto Warburg in 1923^55^ have gained considerable popularity in the past 1.5 decades.^12,56^ Despite its relative popularity, PCLS technology has several drawbacks. First, functionality of PCLS is limited to 3-6 days and even that limited function is ensured by sophisticated perfusion or rocking protocols. Second, the footprint of a typical liver slice is fairly large (~0.5 × 1 cm^2^) which is necessitated by the specifications of microtomes or slicers.^57^ Therefore, slicing and imbedding is not practicable for small wedges, biopsies or fine needle aspirates. As highlighted in this paper, our microfluidic organotypic culture platform addresses the limitations of PCLS technology. It enables 30 days of hepatic function while using minimal amounts of liver tissue. We surveyed PCLS literature for reports of quantifiable functional outputs that can be compared against our rat liver cultures. Values for production of albumin, urea and bile acids for published organotypic rat liver cultures were compiled in Table 1. The levels of these hepatic indicators were standardized using reported experimental details to make comparison across studies possible. We should note that the organotypic liver culture approaches reported to date employed either continuous perfusion or rocking to enhance delivery of nutrients or oxygen. Despite this fact, these existing technologies allowed for 2-6 days of culture while eliciting low levels of hepatic function.^20,21,58,59^ In comparison, our organotypic cultures were 7- to 86-fold smaller, were cultured under simple (static) conditions and remained functional for over 31 days. It is worth pointing out that levels of albumin and urea in our devices at day 25 were significantly higher than those reported for PCLS cultures at much earlier time points (day 4 to 6).

**Table 1:** Comparing functionality of organotypic liver cultures reported to date.

| Author and Year | Culture duration | Sample format | Sample size (mm<sup>3</sup>) | Culture platform | Static/perfused | Albumin (µg/1x10<sup>4</sup>/ml/day) | Urea (µg/1x10<sup>4</sup>/ml/day) | Bile (µM/1x10<sup>4</sup>/ml/day) |
| --- | --- | --- | --- | --- | --- | --- | --- | --- |
| Khong et al. 2007<sup>21</sup> | 3 days | PCLS | 50.27 | Intra-tissue perfusion (ITP) system | Perfused 1mL/min | 0.72 | 2.40 | N/R |
| Hattersley et al. 2008<sup>20</sup> | 2 days | Tissue cube | 4 | Microfluidic device | Perfused 2µL/min | 0.26 x 10<sup>-6</sup> | 1.7 x 10<sup>-2</sup> | N/R |
| Hattersley et al. 2011<sup>58</sup> | 4 days | Tissue cube | 6.28 | Microfluidic device | Perfused 2µL/min | 0.0033 x 10<sup>-6</sup> | 0.64 x 10<sup>-2</sup> | N/R |
| Paish et al. 2019<sup>59</sup> | 6 days | PCLS | 12.56 (x2) | BioPlate | Rocking | 0.53 x 10<sup>-6</sup> | N/R | N/R |
| **Our device** | 31 days | Needle core | 0.58 | Microfluidic device | Static | 1.10 | 3.38 | 1.86 |

### Conclusions

We report the development of microfluidic organotypic liver cultures for long-term maintenance of hepatic function. Our microfluidic device included several features that enabled long-term maintenance of liver tissue: 1) an injection port for simple transfer of liver tissue cores from the needle into a culture chamber, 2) a normally closed valve which could be resealed after tissue injection and allowed to culture microfluidic devices akin to Petri dishes and 3) a design of the culture chamber and transport channels to ensure both sufficient delivery of nutrients and accumulation of hepato-inductive endogenous signals in absence of convection. Microfluidic organotypic rat liver cultures maintained high levels of albumin and urea production over the course of 31 days. These cultures expressed phase I and phase II enzymes and, at day 7, exhibited polarization reminiscent of freshly isolated rat liver tissue. For comparison, phenotype and function declined precipitously in cultures of similarly sized liver tissue in a 96-well plate format over the course of 7 days. Microfluidic liver cultures had 250-fold higher albumin levels compared to conventional well-based culture format at day 7. More impressively, similarly high levels of albumin production persisted in microfluidic cultures until the end of experiment at day 31 of culture. To the best of our knowledge, there have not been reports of comparable duration or functionality for organotypic liver cultures. Beyond demonstrating utility of rodent organotypic liver cultures, we also applied our novel technology to cultivation of human liver tissue. We demonstrate 7 days of function for patient liver tissue in microfluidic devices.

Our microfluidic platform represents a significant step forward for organotypic liver cultures. It ensures long-term phenotype stability and opens new avenues for disease modeling. Minimal tissue requirements afforded by the microfluidic format makes it possible to leverage tissue available from biopsies or fine needle aspirates for personalized therapy testing. Moving forward, we see several areas for further enhancement of the microfluidic organotypic cultures. While current culture conditions (e.g. media) maximize hepatic function, they appear to be suboptimal for maintaining some of the nonparenchymal cell types, namely sinusoidal endothelial cells. We envision refining media composition in the future to address this. Furthermore, we see exciting opportunities to interface microfluidic organotypic cultures with on-chip bioassays for assessing hepatic function and development of integrated therapy-testing platforms.

## Acknowledgements

The authors acknowledge financial support from NIH (DK107255). Additional support was provided by Center of Signaling in Gastroenterology (P30DK084567).

## Conflict of Interest

The authors have no conflicts to disclose.

## Author contributions

J.M.D.H.-V. and A.R. conceived and designed research; J.M.D.H.-V. designed and fabricated the microfluidic device. J.M.D.H.-V. and H.J.H. performed experiments; J.M.D.H.-V., H.J.H., and K.L. analyzed data; J.M.D.H.-V., H.J.H. and G.S. interpreted results of experiments; J.M.D.H.-V. prepared figures; J.M.D.H.-V. drafted manuscript; J.M.D.H.-V. and A.R. edited and revised manuscript. J.M.D.H.- V., H.J.H., K.L., G.S., and A.R. approved final version of manuscript.

## Notes

### Competing Interest Statement

The authors have declared no competing interest.

### Summary of Updates

Supplemental files updated

