## Supplemental Information for "A MICROFLUIDIC DEVICE FOR LONG-TERM MAINTENANCE OF ORGANOTYPIC LIVER CULTURES"

### Supplementary Information

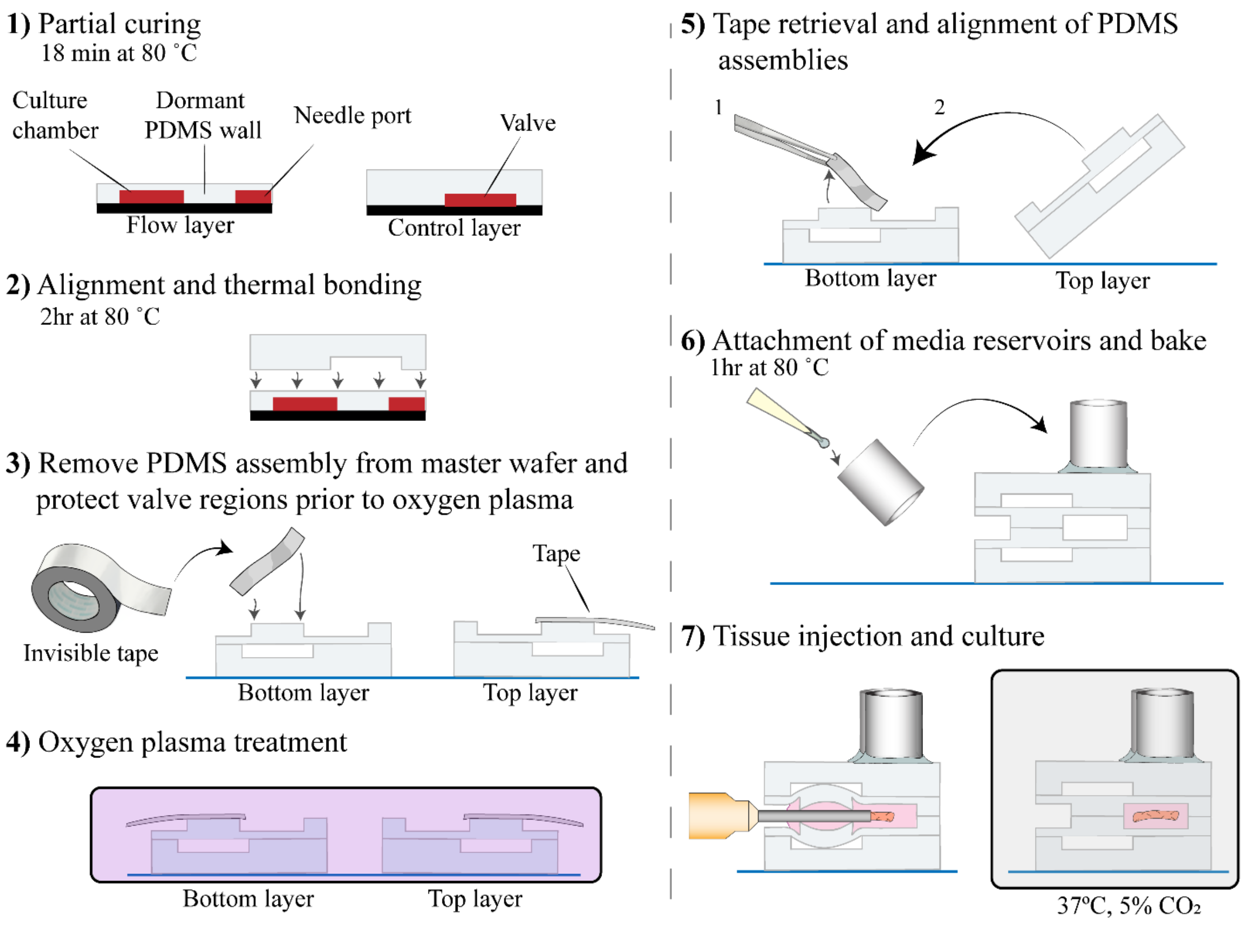


**Figure S1**: **Fabrication of microfluidic devices for culturing liver tissue**. **1**) Replicating photoresist patterns in PDMS to create flow and valve layers of the device. **2**) Thermal bonding of flow and valve layers to create half of the device. **3**) PDMS slab is peeled off from a wafer. The side containing injection port is protected by tape during oxygen plasma treatment. Two halves (mirror images) of the device are treated with oxygen plasma. **4**) devices are aligned and bonded. Regions that were protected during oxygen plasma in the preceding step do not bond to each and may be opened and closed by valve actuation. 5) Cloning cylinders are attached to create media reservoirs. 6) Injection port may be opened or closed by application of vacuum to a valve. This valve is normally closed and devices may be kept in an incubator without connections or vacuum lines.

**Table S1.** Sequences of primers used for RT-PCR analysis.

| Gene | Forward | Reverse |
|---|---|---|
| GAPDH | AGACAGCCGGATCTTCTTGT | CTTGCCGTGGGTAGAGTCAT |
| ALB | CATCCTGAACCGTCTGTGTG | TTTCCACCAAGGACCCACTA |
| HGF | CTTCTGCCGGTCCTGTTG | TCTTCTCTTCTTCTGTCCTTCT |
| TGFβ | CCTGGAAAGGGCTCAACAC | CAGTTCTTCTCTGTGGAGCTG |
| α-SMA | TGCCATGTATGTGGCTATTCA | ACCAGTTGTACGTCCAGAAGC |
| PTPRC | GCTATAAAAAGACCCCTTCAG | CATAGGCAAATAGAGACACTG |
| GFAP | CAAGATGAAACCAACCTGAG | CTTCCTCCTCATGGATCTTC |
| CYP7 | ACGCACCTCGCTATTCTCTG | GGCAGGTCATTCAGTTGCAC |
| CYP2E1 | CCTACATGGATGCTGTGGTG | CTGGAAACTCATGGCTGTCA |
| CYP1A2 | ACCATCCCCCACAGTACAA | GTTGACCTGCCACTGGTTTA |
| CYP1A1 | TGAGTTTGGGGAGGTTACTGGTT | TGAAGGCATCCAGGGAAGAGT |
| CD163 | ATAGTCTGCTCACGATACATAG | GACATAAGACATGAGCATCTTC |
| PECAM1 | AAAACCACAATTGAGTACCAG | ACTTAGCTTGACGTTCTTTG |
| DESMIN | AAAGAAGAGAGAGGCAGAG | TCTTTATTGTTTCTGTCCAGG |
| GSTM1 | AGCTCATCATGCTTTGTTAC | AGTAGAGCTTCATCTTCTCAG |
| UGT1A1 | CTTTGTGAAAGATTACCCCAG | GACATAGGCTTCAAATTCCTG |

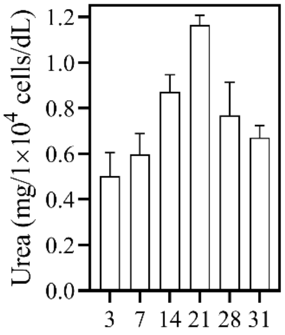


**Figure S2**: Long-term maintenance of urea synthesis in microfluidic organotypic cultures.

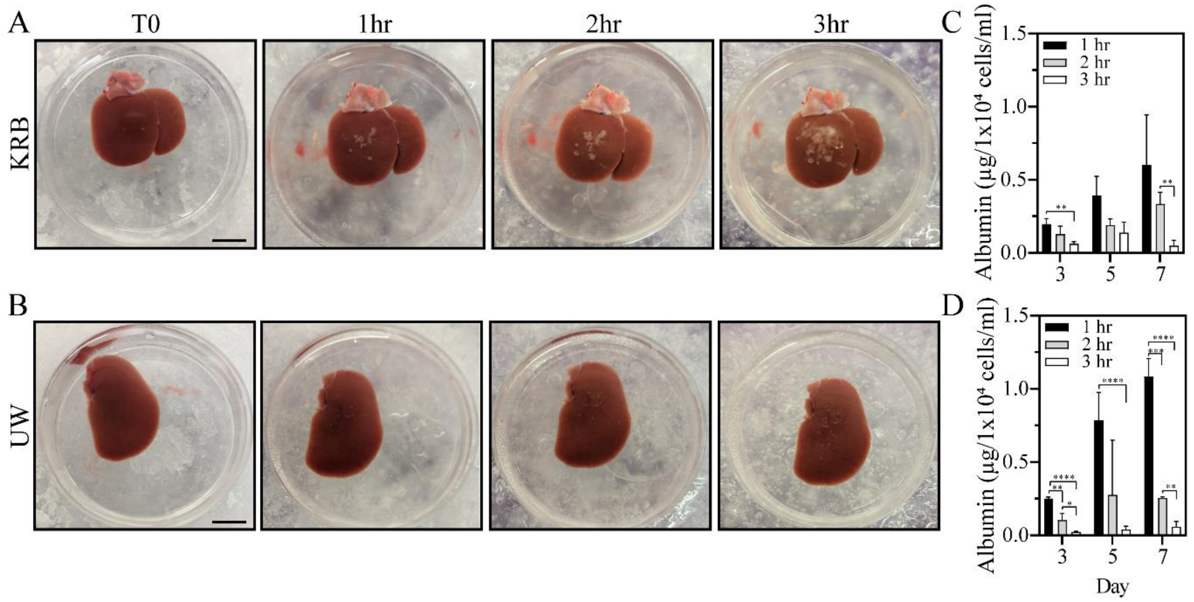


**Figure S3**: **Effects of preservation solution and storage time on functionality of microfluidic liver cultures**. (**A**) Left liver lobe preserved in KRB solution. (**B**) Right liver lobe was preserved in UW cold storage solution. Liver lobes were preserved for 3 hr and needle core were collected every hour and cultured in the microfluidic devices for 7 days. (**C**) Albumin production for liver tissue preserved in KRB for different duration of time prior to microfluidic cultures. (**D**) Albumin synthesis for liver tissue preserved in UW solution prior to microfluidic cultures. Please provide details about statistical analysis (samples, number of dots and statistical significance).  Data are represented as means ±SD of 3 tissue samples cultured in well and microfluidic platform. Statistical significance determined by Mann–Whitney test, * p-value <0.05, ** p-value <0.01, *** p-value < 0.001, and **** p-value < 0.0001.

**Video 1: Introduction of a liver needle core into a microfluidic device.**

**Video 2: Injection of a second needle core into a microfluidic device for coculture purposes.**
